# Pre-Hospital Midazolam for Treatment of Status Epilepticus Before and After the Rapid Anticonvulsant Medication Prior to Arrival Trial: A National Observational Cohort Study

**DOI:** 10.1101/084327

**Authors:** Eytan Shtull-Leber, Robert Silbergleit, William J. Meurer

## Abstract

**Background:** Implementation of evidence-based treatment for pre-hospital status epilepticus can improve outcomes. We hypothesized that publication of a pivotal pre-hospital clinical trial (RAMPART), demonstrating superiority of intramuscular midazolam over intravenous lorazepam, altered the national utilization rates of midazolam for pre-hospital treatment of status epilepticus, while upholding its safety and efficacy outside the trial setting.

**Methods and Findings:** This is a retrospective, observational cohort study of pre-hospital patient encounters throughout the United States in the National Emergency Medicine Services Information System database, from January 2010 through December 2014. We compared the rates and odds of midazolam use as the first-line treatment for status epilepticus among all adult and pediatric benzodiazepine-treated seizures before and after RAMPART publication (February 2012). Secondary analyses were conducted for rates of airway interventions and rescue therapy, as proxies for safety and efficacy of seizure termination. 156,539 benzodiazepine-treated seizures were identified. Midazolam use increased from 26.1% in January 2010 to 61.7% in December 2014 (difference +35.6%, 95% CI, 32.7%-38.4%). The annual rate of midazolam adoption increased significantly from 5.9% per year to 8.9% per year after the publication of RAMPART (difference +3.0% per year; 95%CI, 1.6%-4.5% per year; adjusted OR 1.24; 95%CI, 1.17-1.32). Overall frequency of rescue therapy and airway interventions changed little after the publication of RAMPART.

**Conclusions:** These data are consistent with effective, ongoing, but incomplete clinical translation of the RAMPART results. The effects of the trial, however, cannot be isolated. The safety and effectiveness of midazolam for treatment of seizures in prehospital clinical practice appear consistent with trial data, which should encourage continuing increases in utilization.

## Main Manuscript

### Abbreviations

95% Cl denotes 95% confidence interval, ALS advanced life support, BLS basic life support, EMS emergency medical services, ICC, intra-class correlation, IV intra-venous, NEMSIS National EMS Information System, OR odds ration, RAMPART Rapid Anticonvulsant Medication Prior to Arrival.

### Introduction

Second state knowledge translation—the process by which clinical trial findings move into accepted clinical practice—has often been slow and incomplete. Indeed, the Institute of Medicine reported an average of 17 years interval between reports of randomized controlled trials and broad incorporation into practice[1]. Very little is known about knowledge translation in the pre-hospital setting, where a historical dearth of robust clinical trials and the multiple levels of medical providers may be barriers to change in practice[2].

The 2012 publication of the results of the Rapid Anticonvulsant Medication Prior to Arrival Trial (RAMPART) [3] provides a unique opportunity to examine the effect of a high-impact factor journal on second stage knowledge translation in the pre-hospital environment. RAMPART was a large multicenter randomized clinical trial of pre-hospital treatment of patients with status epilepticus. The trial demonstrated that intramuscular midazolam, which was increasingly being used based on theoretical advantages, was superior to intravenous lorazepam at terminating convulsions prior to emergency department arrival without the need for rescue medication.

We therefore determined the changes in the pre-hospital use of midazolam for the treatment of status epilepticus before and after publication of RAMPART. Additionally, we evaluated the generalizability of the RAMPART findings by evaluating the effectiveness of pre-hospital midazolam for the termination of seizures in routine clinical practice on a national level.

## Methods

### Overview and setting

This is a phase IV, retrospective, observational cohort study of the impact of the publication of the RAMPART results on the use of midazolam for the treatment status epilepticus in the prehospital setting. The RAMPART trial began in June 2009, concluded in January 2011, and the results were published in the New England Journal of Medicine in February 2012. The University of Michigan Institutional Review Board determined this project, which used de-identified data, was not regulated human subjects research.

### Data source

The National Emergency Medical Services Information System (NEMSIS) is a national repository of standardized patient care reports from EMS activations from up to 48 U.S. States and territories depending on year.[4] Data represent a convenience sample collected by EMS agencies, submitted to state repositories, and consolidated into the NEMSIS registry after a validation and data cleaning process[4].

Using the NEMSIS dataset, we identified all patient care events involving adult and pediatric patients who were treated with benzodiazepines for presumed status epilepticus in the prehospital setting, from January 1, 2010, through December 31, 2014. As there is no diagnostic code to differentiate status epilepticus from other seizures, status epilepticus was operationalized as the presence of a seizure and the administration of a benzodiazepine. We utilized two sampling strategies for seizures with the following Field Value codes. First, events were sampled where seizure was coded in either the chief complaint as reported by dispatch (455), the CMS condition code (8017), or the EMS provider’s primary (1710) or secondary impression (1845). Second, sampling was narrowed to the primary or secondary impression. After limiting the samples to those treated with benzodiazepines, the total eligible population was similar for the two seizure sampling strategies, so the latter, narrow strategy was chosen for a more precise operational definition of status epilepticus.

### Measures

The primary outcome measure is midazolam use as the first-line treatment for status epilepticus, as a proportion of all benzodiazepine-treated seizures, evaluated monthly. First-line medication was defined as the first medication listed in the “medications given” variable for a patient care event. While RAMPART demonstrated the efficacy specifically of intramuscular midazolam, medication route is not available in the NEMSIS database. Venous access procedures could not be used as a proxy for intravenous (IV) administration of medications, as they were equally as common for those treated with midazolam as for those treated with other, IV-administrable, benzodiazepines (67.0% and 68.8%, respectively).

The main predictor is an indicator variable for events occurring after RAMPART publication in February 2012. Age, reported in units of hours, days, months, and years, was converted to years and analyzed continuously and categorically according to the age categories used in RAMPART. Gender was evenly distributed at baseline (50.8% male, 49.2% female).

Patient race (black, white, other) and ethnicity (Hispanic, not Hispanic) were included to look for disparities in rates of knowledge translation. Geographic distribution of events was defined by US Census Regions (Northeast, Midwest, South, West), and levels of urbanicity (urban, suburban, rural, wilderness). Characteristics of the EMS encounter, including service level (basic life support (BLS), advanced life support (ALS), air, other), primary role of the unit (transport, other), and type of service requested (911 response, inter-facility transfer, other) were included given our a priori belief of an association with midazolam use. Average annual call volume for represented years was calculated and used as a proxy for EMS agency size.

Secondary outcome measures include airway intervention and rescue therapy, as proxies for safety and efficacy of seizure termination, respectively. Airway intervention was defined as any invasive procedure to secure the tracheal airway, including any of the following procedure codes: 31.420 (direct laryngoscopy), 96.042 (rapid sequence intubation), 96.051 (combitube blind insertion airway device), 96.030 (esophageal obturator airway/esophageal gastric tube airway), 96.053 (King LT blind insertion airway device), 96.052 (laryngeal mask blind insertion airway device), 96.041 (nasotracheal intubation), 96.040 (orotracheal intubation), and 31.120 (surgical cricothyrotomy). Rescue therapy, or the administration of a second drug or dose to terminate a seizure, was defined by the presence of a second benzodiazepine listed under “medications given.”

Explanatory variables for secondary analysis are identical to the primary analysis with the addition of a binary variable for the use of midazolam versus other benzodiazepines, and its interaction with time.

### Statistical analysis – predefined

Midazolam use as a proportion of all benzodiazepines was plotted graphically by month, and by levels of each categorical predictor, covering two phases: 1) pre-RAMPART publication phase (January 2010 through January 2012), and 2) post-RAMPART publication phase (February 2012 through December 2014). Rates of midazolam adoption during the two phases were compared using a piecewise linear regression model (spline).

A series of nested random-effects hierarchical logistic regression models, using Laplace approximation, were fit with a primary outcome variable of midazolam use versus all other benzodiazepines. A random intercept for EMS agency was applied to adjust for and measure within-agency correlation. Intra-class correlation (ICC) was calculated using the method of Snijders and Bosker (Equation below) [5]. The numerator is the level 2 (EMS agency) variance and the denominator is the level 2 variance plus the standard logistic distribution variance (level 1).

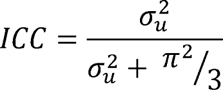

Model 0 is the intercept-only model, which assesses the proportion of midazolam used by the average EMS agency during the study period. Model 1 assesses the unadjusted impact of RAMPART publication. Model 2 assesses the impact of RAMPART publication, adjusting for secular trends. Model 3 further adjusts for patient demographics. Model 4, the fully adjusted model, further adjusts for characteristics of the EMS call.

Three variables (race, ethnicity, and service level) were omitted from the logistic regression models due to high percentages of missing data, which would significantly limit the number of observations in the more fully adjusted models. All other variables had less than 1.5% missing data and were included in the models. In order to reduce any bias caused by diminishing numbers of events in later models that incorporate variables with missing data, all nested models were limited to events with non-missing data for all variables in the final model. Sensitivity analyses were carried out with the inclusion of each of the 3 highly missing variables included in the fully adjusted model. The census region Island Areas was omitted from regression analysis due to low representation.

Secondary analyses compared rates of airway interventions and rescue therapy before and after RAMPART publication and between those treated with midazolam and other benzodiazepines. Because midazolam dose and route may have also changed over time or in response to the trial, we also modeled the interaction with time for each agent in a series of nested hierarchical logistic regression models for the prediction of secondary outcomes. Model 0 is the intercept-only models, which assesses the frequency of rescue therapy and airway interventions by the average EMS agency during the study period. Model 1 assesses the unadjusted impact of midazolam use. Model 2 assesses the unadjusted interaction of midazolam use and time. Model 3 assesses the interaction of midazolam use and time, adjusted for demographics. Model 4, the fully adjusted models, further adjust for characteristics of the EMS call.

We conducted data manipulation and analysis with SAS (v. 9.4, SAS Institute, Cary, NC).

### Statistical analysis – post hoc

After observing the rate of midazolam use appeared to increase more rapidly at the end of 2011, prior to the publication of RAMPART, we conducted two post-hoc analyses of potential confounders on the rate of change in midazolam adoption. A spline model of midazolam use among EMS agencies that contributed data to NEMSIS in all 5 study years was used to reduce artifact that could reflect the onboarding of new states and EMS agencies into the dataset at the beginning of each year. Additionally, we examined the effect of the national diazepam shortages on benzodiazepine use patterns with a series of spline models with indicator variables for the months in which national diazepam shortages began, were publically announced, and subsequently resolved. Effects of lorazepam shortages were not investigated as the diversity of manufacturers was felt to likely preclude any perceptible impact.

After fitting the hierarchical logistic models of the primary outcome and seeing the high ICC, we fit a multinomial logistic regression model of midazolam use levels per EMS agency-year (none, some, most, all) in order to determine the effect of agency-level characteristics on the overall odds of midazolam use. We chose multinomial logistic regression despite the ordered nature of the data as we could not dependably assume equal odds associations between midazolam use levels. In addition, the high proportion of agencies using no midazolam or all midazolam made Poisson and linear regression distributionally inappropriate. Predictors included the proportion of each agency’s patient population that is in a particular age bracket (<5 years old, 6-20 years old, ≥60 years old), or is female, black, or Hispanic. Additional predictors included agency size (annual call volume, size quintile), catchment area (number of counties in which agency operates), urbanicity (defined by > 95% of encounters occurring in a single urbanicity level, otherwise mixed), highest service level (specialty care only, BLS, ALS, Air), transport capability, and provision of 911 scene response. The model was limited to agency-years for which each of the above variables was nonmissing. There were no problems with multicollinearity, with a variance inflation factor threshold of 5.

## Results

Among the 93.7 million patient care events (representing 10,830 EMS agencies) in the NEMSIS database from 2010-2014 (Fig 1), 1.9 million were characterized as a seizure by the provider’s primary or secondary impression. Of those, 156,539 (from 3,504 EMS agencies) were treated with a known benzodiazepine. All multivariable regression models analyzed the 152,662 cases without any missing data for explanatory variables that were included in the final model.

**Fig 1.**
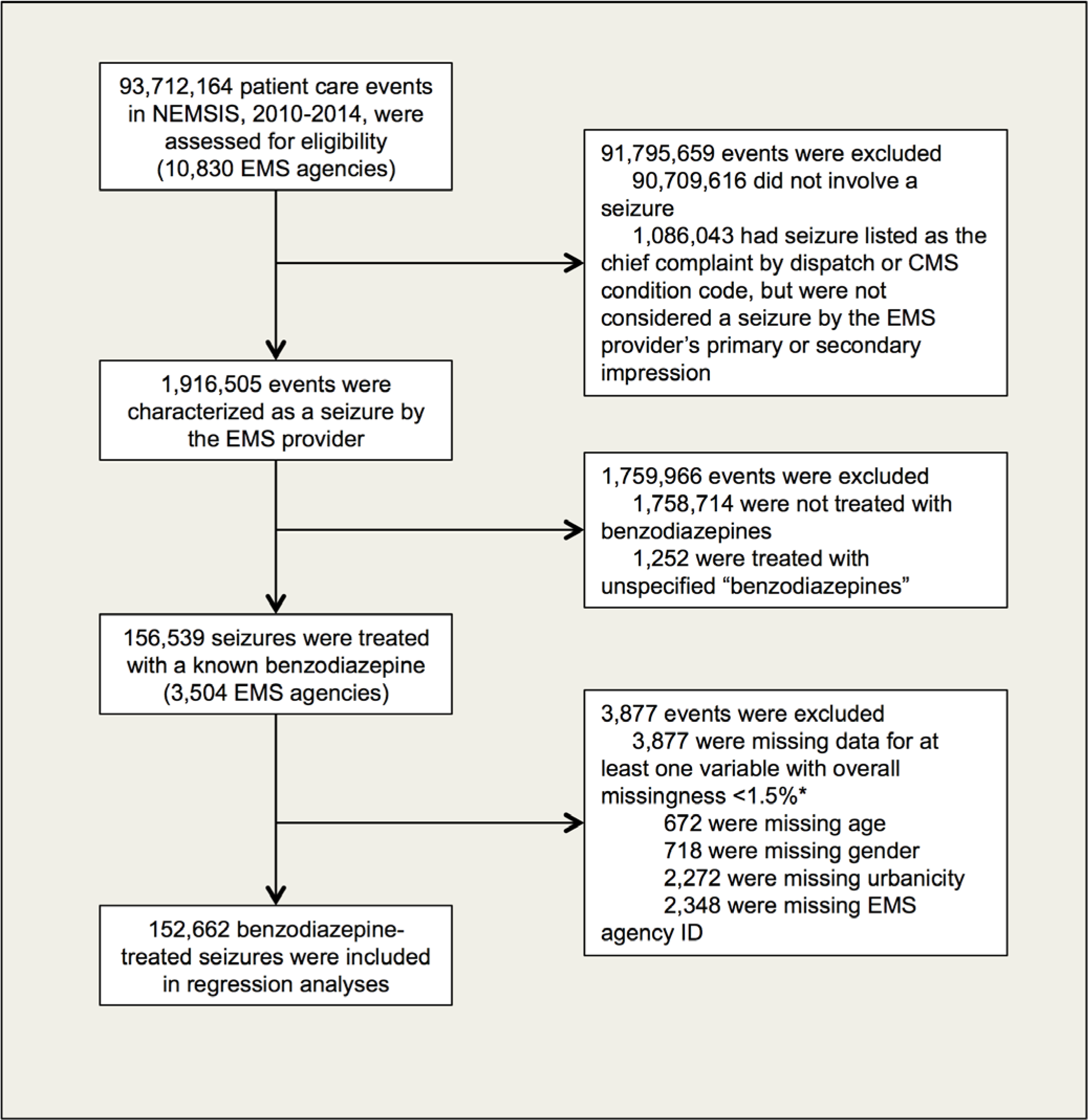
Inclusion of eligible patient care events for primary analysis. *Race, ethnicity and service level all had high levels of missingness and were therefore excluded from the regression models in order to maximize the number of events analyzed. Race was missing for *27,770* events, ethnicity for 43,164 events, and service level for 66,683 events.

The mean patient age was 39.4 ± 21.8 years (median 39, interquartile range 24-55 years). Additional event characteristics for each year of the study are listed in Table 1, by categorical predictors. Age is reported by categories due to lack of a linear trend. Census region, urbanicity, service level, primary role of unit, and service type all appear to be linearly associated with time. Table 2 lists the number of seizures that were treated with midazolam in each year of the study, by categorical predictors. Census region, urbanicity, service level, primary role of unit, and service type also appear to be associated with the use of midazolam and are potential confounders.

**Table 1.** Characteristics of EMS encounters involving status epilepticus in the NEMSIS database, 2010-2014.

| Covariates | 2010, No. (%)<sup>a</sup> | 2011, No. (%) | 2012, No. (%) | 2013, No. (%) | 2014, No. (%) |
| --- | --- | --- | --- | --- | --- |
| Patient age | (n=16.2)<sup>b</sup> | (n=23.7) | (n=32.2) | (n=39.6) | (n=44.2) |
| 0-5 Years | 1.5 (9.3) | 2.2 (9.2) | 2.9 (8.6) | 3.32 (8.3) | 3.6 (8.1) |
| 6-10 Years | 0.5 (3.2) | 0.7 (3.2) | 1.0 (3.0) | 1.2 (2.9) | 1.4 (3.1) |
| 11-20 Years | 1.5 (9.2) | 2.1 (8.9) | 2.8 (8.7) | 3.54 (8.8) | 3.9 (8.8) |
| 21-40 Years | 5.0 (30.8) | 7.3 (30.9) | 10.1 (31.5) | 12.5 (31.5) | 14.0 (31.7) |
| 41-60 Years | 5.0 (31.0) | 7.3 (30.9) | 10.0 (31.0) | 12.4 (31.2) | 13.6 (30.7) |
| ≥ 60 Years | 2.6 (16.4) | 4.0 (17.0) | 5.4 (16.8) | 6.8 (17.3) | 7.8 (17.6) |
| Patient gender | (n=16.3) | (n=23.7) | (n=32.2) | (n=39.6) | (n=44.0) |
| Male | 8.3 (50.9) | 12.1 (50.9) | 16.4 (51.1) | 19.9 (50.4) | 22.4 (50.8) |
| Female | 8.0 (49.1) | 11.7 (49.2) | 15.8 (49.0) | 19.6 (49.6) | 21.7 (49.2) |
| Patient race | (n=13.5) | (n=20.0) | (n=26.8) | (n=32.7) | (n=35.8) |
| White | 8.8 (65.5) | 13.0 (65.0) | 17.8 (66.6) | 21.32 (65.0) | 22.9 (64.0) |
| Black | 3.4 (25.6) | 5.6 (27.9) | 7.1 (26.3) | 8.9 (27.3) | 10.0 (27.9) |
| Other | 1.2 (8.8) | 1.4 (7.1) | 1.9 (7.1) | 2.5 (7.7) | 2.9 (8.1) |
| Patient ethnicity | (n=11.6) | (n=17.2) | (n=23.4) | (n=29.5) | (n=31.8) |
| Hispanic | 1.0 (8.4) | 1.3 (7.4) | 1.7 (7.3) | 2.4 (8.3) | 2.7 (8.4) |
| Not Hispanic | 10.6 (91.6) | 15.9 (92.6) | 21.7 (92.7) | 27.1 (91.7) | 29.1 (91.6) |
| Region | (n=16.4) | (n=23.8) | (n=32.3) | (n=39.7) | (n=44.3) |
| Northeast | 1.5 (9.4) | 2.8 (11.7) | 4.5 (14.0) | 5.1 (12.8) | 6.6 (14.8) |
| Midwest | 3.5 (21.4) | 4.6 (19.5) | 7.0 (21.7) | 7.4 (18.6) | 9.2 (20.7) |
| South | 9.0 (55.2) | 13.9 (58.5) | 17.3 (53.4) | 20.5 (51.7) | 20.9 (47.3) |
| West | 2.3 (13.9) | 2.4 (10.1) | 3.5 (10.8) | 6.7 (16.9) | 7.6 (17.2) |
| Island areas | 0.02 (0.1) | 0.03 (0.1) | 0.03 (0.1) | 0.04 (0.1) | 0.05 (0.1) |
| Urbanicity | (n=16.3) | (n=23.7) | (n=31.8) | (n=39.2) | (n=43.3) |
| Urban | 12.2 (74.9) | 18.0 (75.9) | 24.5 (77.1) | 31.3 (79.9) | 35.0 (80.8) |
| Suburban | 1.6 (9.8) | 2.0 (8.5) | 2.7 (8.5) | 3.1 (8.0) | 3.3 (7.7) |
| Rural | 2.0 (12.6) | 3.0 (12.9) | 3.8 (12.0) | 3.8 (9.7) | 4.0 (9.3) |
| Wilderness | 0.4 (2.8) | 0.6 (2.7) | 0.8 (2.5) | 0.9 (2.3) | 0.9 (2.2) |
| Service level | (n=9.8) | (n=13.9) | (n=18.4) | (n=22.5) | (n=25.2) |
| BLS<sup>c</sup> | 0.3 (3.0) | 0.32 (1.9) | 0.3 (1.8) | 0.3 (1.3) | 0.3 (1.3) |
| ALS<sup>c</sup> | 9.1 (93.7) | 13.1 (94.1) | 17.2 (93.2) | 21.1 (94.0) | 23.5 (93.3) |
| Air | 0.3 (2.9) | 0.5 (3.4) | 0.8 (4.4) | 0.9 (4.1) | 1.1 (4.3) |
| Other | 0.03 (0.4) | 0.09 (0.6) | 0.1 (0.6) | 0.1 (0.6) | 0.3 (1.1) |
| Unit role | (n=16.4) | (n=23.8) | (n=32.3) | (n=39.7) | (n=44.3) |
| Transport | 15.4 (94.3) | 22.0 (92.5) | 29.9 (92.6) | 37.0 (93.1) | 40.8 (92.0) |
| Other | 0.9 (5.7) | 1.8 (7.5) | 2.4 (7.4) | 2.7 (6.9) | 3.6 (8.0) |
| Service type | (n=16.4) | (n=23.8) | (n=32.3) | (n=39.7) | (n=44.3) |
| 911 Response | 15.3 (93.3) | 22.3 (93.7) | 30.3 (93.7) | 37.2 (93.7) | 41.4 (93.3) |
| Transfer | 0.6 (3.5) | 0.98 (3.7) | 1.3 (4.0) | 1.6 (4.0) | 1.9 (4.2) |
| Other | 0.5 (3.2) | 0.6 (2.7) | 0.8 (2.4) | 1.0 (2.4) | 1.1 (2.5) |
| Agency size quintile | (n=16.2) | (n=23.7) | (n=31.7) | (n=39.1) | (n=43.4) |
| 1st | 0.2 (1.3) | 0.4 (1.6) | 0.5 (1.6) | 0.6 (1.6) | 0.8 (1.7) |
| 2nd | 0.5 (3.3) | 0.9 (3.7) | 1.2 (3.7) | 1.4 (3.7) | 1.7 (3.9) |
| 3rd | 1.0 (6.2) | 1.7 (7.2) | 2.5 (8.0) | 2.8 (7.0) | 3.2 (7.3) |
| 4th | 2.2 (13.8) | 3.4 (14.2) | 4.9 (15.6) | 5.5 (14.1) | 5.8 (13.3) |
| 5th | 12.2 (75.4) | 17.4 (73.4) | 22.5 (71.1) | 28.8 (73.6) | 32.1 (73.8) |

<sup>a</sup>Number of events, in thousands (% of non-missing values for the variable in a given year)
<sup>b</sup>Number of events varied by covariate and year based on number of missing values in NEMESIS.
Across all years, number of missing values were as follows: age 672 (0.4%); gender 718 (0.5%); race 27,770 (17.7%); ethnicity 43,164 (27.6%); urbanicity 2,272 (1.5%); CMS service level 66,683 (42.6%); Agency ID, which was used to create agency size quintiles, 2,348 (1.5%). There were no missing data for census region, primary role of unit, or service type.
<sup>c</sup>BLS = basic life support, ALS = advanced life support.

**Table 2.** Number of pre-hospital cases of status epilepticus treated with Midazolam (% of all benzodiazepine-treated seizures), 2010-2014.

| Covariates | 2010,<br>No. (%)<sup>a</sup> | 2011,<br>No. (%) | 2012,<br>No. (%) | 2013,<br>No. (%) | 2014,<br>No. (%) | Difference in<br>proportions, (95%<br>CI) |
| --- | --- | --- | --- | --- | --- | --- |
| Total | 45.8 (28.0) | 71.7 (30.1) | 133.8 (41.4) | 197.3 (49.7) | 252.0 (56.9) | 28.9 (28.0-29.7) |
| Patient age |  |  |  |  |  |  |
| 0-5 Years | 4.2 (28.1) | 7.1 (32.7) | 12.0 (41.7) | 16.7 (50.9) | 21.0 (58.7) | 30.6 (27.8 to 33.3) |
| 6-10 Years | 1.3 (25.2) | 1.9 (25.6) | 3.9 (39.9) | 5.3 (46.0) | 7.4 (54.8) | 29.6 (24.8 to 34.0) |
| 11-20 Years | 4.3 (28.9) | 6.5 (30.6) | 11.3 (40.4) | 16.2 (46.5) | 21.5 (55.5) | 26.6 (23.8 to 29.4) |
| 21-40 Years | 14.3 (28.6) | 23.5 (32.1) | 42.8 (42.2) | 64.6 (51.8) | 81.1 (57.8) | 29.2 (27.7 to 30.7) |
| 41-60 Years | 14.2 (28.3) | 21.2 (29.0) | 41.5 (41.6) | 61.6 (49.8) | 77.7 (57.3) | 29.0 (27.4 to 30.4) |
| ≥ 60 Years | 7.0 (26.4) | 11.2 (27.8) | 21.6 (39.8) | 32.2 (47.1) | 42.5 (54.6) | 28.3 (26.2 to 30.3) |
| Patient gender |  |  |  |  |  |  |
| Male | 22.5 (27.2) | 36.3 (30.1) | 68.3 (41.5) | 100.0 (50.1) | 127.3 (56.9) | 29.8 (28.6 to 30.9) |
| Female | 23.1 (29.0) | 35.1 (30.1) | 65.0 (41.23) | 96.3 (49.1) | 123.1 (56.8) | 27.9 (26.7 to 29.1) |
| Patient race |  |  |  |  |  |  |
| White | 24.3 (27.6) | 37.0 (28.4) | 67.8 (38.0) | 96.0 (45.1) | 118.8 (51.9) | 24.4 (23.2 to 25.5) |
| Black | 8.8 (25.6) | 16.6 (29.7) | 30.8 (43.6) | 44.9 (50.1) | 58.4 (58.6) | 33.0 (31.2 to 34.7) |
| Other | 3.8 (32.0) | 4.7 (32.9) | 7.8 (41.2) | 13.0 (52.0) | 18.1 (62.2) | 30.2 (27.0 to 33.3) |
| Patient ethnicity |  |  |  |  |  |  |
| Hispanic | 2.7 (28.0) | 3.8 (29.8) | 6.9 (40.2) | 12.6 (51.7) | 16.0 (59.7) | 31.7 (28.2 to 35.0) |
| Not Hispanic | 28.7 (27.1) | 44.7 (28.1) | 84.3 (38.9) | 127.6 (47.1) | 155.9 (53.6) | 26.5 (25.5 to 27.5) |
| Region |  |  |  |  |  |  |
| Northeast | 2.9 (18.9) | 5.4 (19.2) | 16.2 (35.8) | 22.1 (43.5) | 33.8 (51.6) | 32.6 (30.3 to 34.9) |
| Midwest | 15.4 (44.1) | 20.6 (44.3) | 35.6 (50.7) | 40.7 (55.1) | 60.4 (66.0) | 21.9 (20.0 to 23.8) |
| South | 21.2 (23.5) | 37.0 (26.6) | 64.8 (37.6) | 90.4 (44.0) | 105.5 (50.4) | 26.9 (25.8 to 28.0) |
| West | 6.3 (27.6) | 8.7 (36.1) | 17.2 (49.3) | 44.1 (65.8) | 52.1 (68.4) | 40.8 (38.7 to 42.9) |
| Island areas | 0.0 (0.0) | 0.0 (0.0) | 0.0 (0.0) | 0.03 (7.5) | 0.1 (21.3) | 21.3 (2.1 to 34.9) |
| Urbanicity |  |  |  |  |  |  |
| Urban | 37.3 (30.5) | 58.9 (32.8) | 109.3 (44.6) | 168.5 (53.8) | 215.6 (61.6) | 31.1 (30.1 to 32.0) |
| Suburban | 3.3 (20.4) | 4.4 (21.8) | 8.2 (30.4) | 10.7 (33.9) | 13.1 (39.1) | 18.7 (16.1 to 21.2) |
| Rural | 4.3 (21.1) | 7.0 (23.1) | 13.2 (34.6) | 13.2 (34.7) | 15.7 (39.0) | 17.9 (15.5 to 20.2) |
| Wilderness | 0.8 (17.0) | 0.8 (13.0) | 1.7 (22.0) | 2.3 (24.9) | 3.3 (34.5) | 17.5 (12.8 to 22.0) |
| Service level |  |  |  |  |  |  |
| BLS<sup>b</sup> | 1.5 (51.7) | 0.7 (27.6) | 1.5 (44.3) | 1.5 (51.5) | 1.8 (52.8) | 1.1 (-6.7 to 8.9) |
| ALS<sup>b</sup> | 22.2 (24.3) | 38.5 (29.4) | 69.9 (40.7) | 99.9 (47.2) | 128.7 (54.7) | 30.4 (29.3 to 31.5) |
| Air | 1.2 (41.4) | 1.9 (40.6) | 3.0 (37.3) | 4.3 (46.7) | 4.5 (41.5) | 0.1 (-6.4 to 6.4) |
| Other | 0.2 (58.8) | 0.3 (36.7) | 0.4 (43.6) | 0.9 (62.4) | 1.3 (44.6) | -14.2 (-3.4 to 30.1) |
| Unit role |  |  |  |  |  |  |
| Transport | 43.2 (28.0) | 67.5 (30.6) | 125.3 (41.9) | 181.5 (49.1) | 228.1 (55.9) | 28.0 (27.1 to 28.8) |
| Other | 264 (28.1) | 4.2 (23.5) | 8.4 (35.4) | 15.7 (57.4) | 23.9 (67.3) | 39.2 (35.9 to 42.4) |
| Service type |  |  |  |  |  |  |
| 911 Response | 42.4 (27.7) | 66.7 (29.9) | 127.0 (41.9) | 186.7 (50.2) | 239.1 (57.8) | 30.1 (29.2 to 30.9) |
| Transfer | 2.2 (37.7) | 3.2 (36.7) | 4.2 (33.0) | 6.2 (39.3) | 7.1 (38.2) | 0.5 (-4.1 to 5.0) |
| Other | 1.3 (24.8) | 1.8 (28.1) | 2.6 (34.3) | 4.4 (46.5) | 5.8 (52.0) | 27.3 (22.4 to 31.9) |
| Agency size quintile |  |  |  |  |  |  |
| 1st | 0.4 (18.1) | 1.1 (29.0) | 2.0 (38.6) | 2.9 (47.3) | 3.7 (49.7) | 31.6 (24.8 to 37.5) |
| 2nd | 1.6 (29.6) | 2.5 (28.6) | 4.9 (41.5) | 7.0 (48.3) | 8.7 (52.1) | 22.5 (17.9 to 26.9) |
| 3rd | 2.8 (28.4) | 4.8 (27.9) | 9.9 (39.0) | 12.8 (46.5) | 16.5 (52.2) | 23.8 (20.4 to 27.0) |
| 4th | 6.8 (30.4) | 10.5 (31.3) | 20.1 (40.8) | 25.2 (45.5) | 29.5 (50.8) | 20.4 (18.1 to 22.7) |
| 5th | 33.3 (27.3) | 52.4 (30.1) | 95.1 (42.2) | 146.6 (50.9) | 189.2 (59.2) | 31.9 (30.9 to 32.8) |

<sup>a</sup>Number of midazolam-treated events, in hundreds, (% of all benzodiazepine-treated events, from corresponding cell in Table 1)
<sup>b</sup>BLS = basic life support, ALS = advanced life support

### Midazolam use

Midazolam use for pre-hospital status epilepticus increased from 26.1% (326 of 1249) to 61.7% (2412 of 3909) of cases, between January 2010 and December 2014 (difference +35.6%; 95% CI, 32.7%-38.4%). This corresponds to a 7.7% increase per year (95% CI, 7.3%-8.1% per year). Midazolam use generally increased over time for all categorical predictors (Fig 2). Spline regression demonstrated a significant increase in the rate of midazolam adoption, from a pre-RAMPART rate of 5.9% per year to a post-RAMPART rate of 8.9% per year (difference +3.0% per year; 95%CI, 1.6%-4.5% per year; Fig 3). Post-hoc regression analyses that limited data to the 1761 (50%) EMS agencies that contributed data to NEMSIS in all 5 study years (Fig 4) and that accounted for the effects of the diazepam shortage (Fig 5) also demonstrated a significant increase in the rate of midazolam adoption after the publication of RAMPART, and may explain the apparent increase in midazolam adoption prior to RAMPART publication.

**Fig 2.**
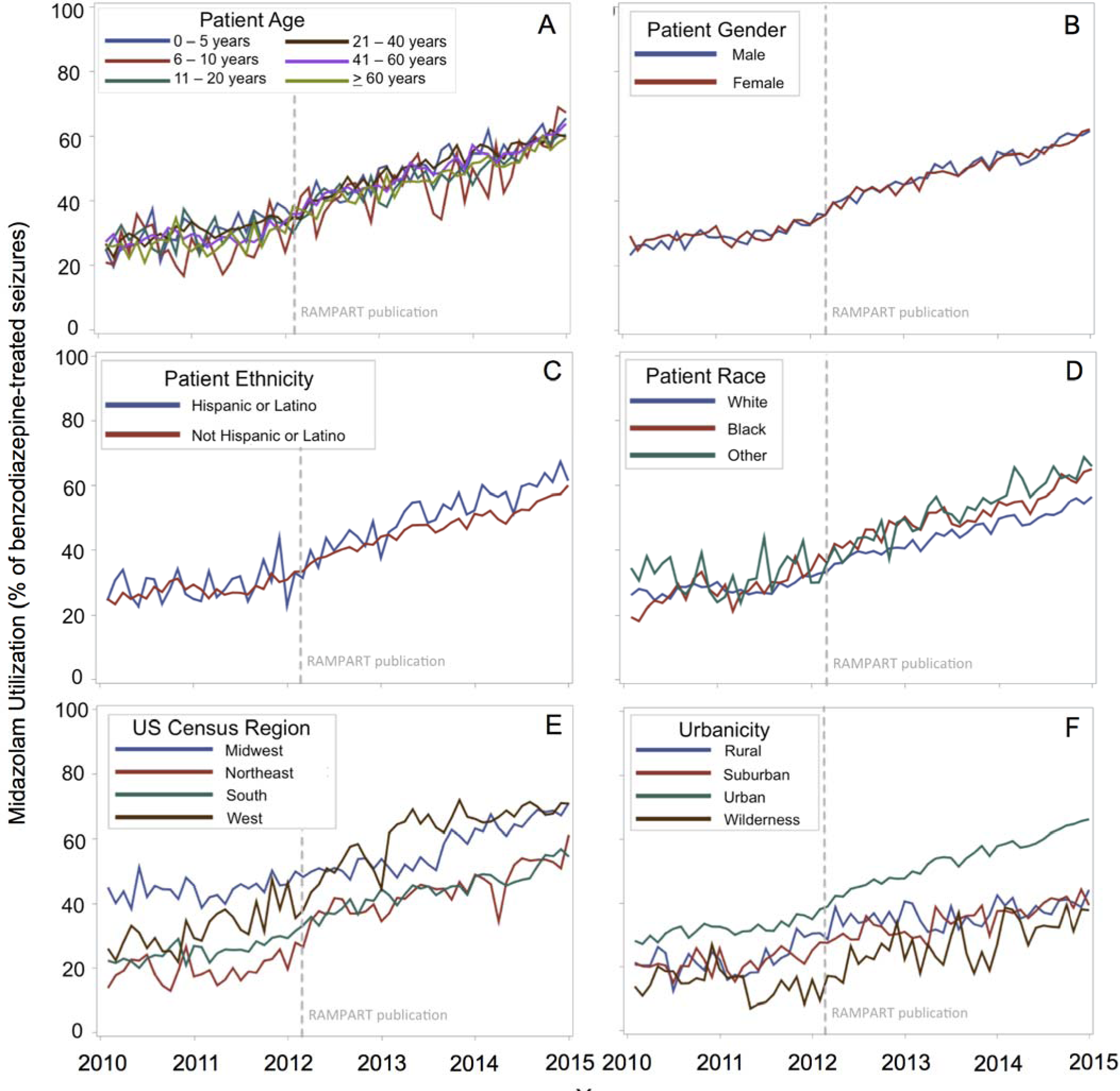
Midazolam utilization for the treatment of seizures in the pre-hospital setting. First-line treatment of status epilepticus with midazolam, as a percentage of all benzodiazepine-treated seizures^a^, evaluated monthly is presented by patient demographic characteristics. Midazolam use generally increased over time for all categorical predictors (A – F), with the greatest variation by geography (E, F). ^a^The denominator varied by covariate and year 249 based on frequency of missing data, with n= 155,867 for age (A), 155,821 for gender (B), 113,375 for ethnicity (C), 128,769 for race (D), 154 for urbanicity (F), and 156,539 for census region, (E) which had no missing data.

**Fig 3.**
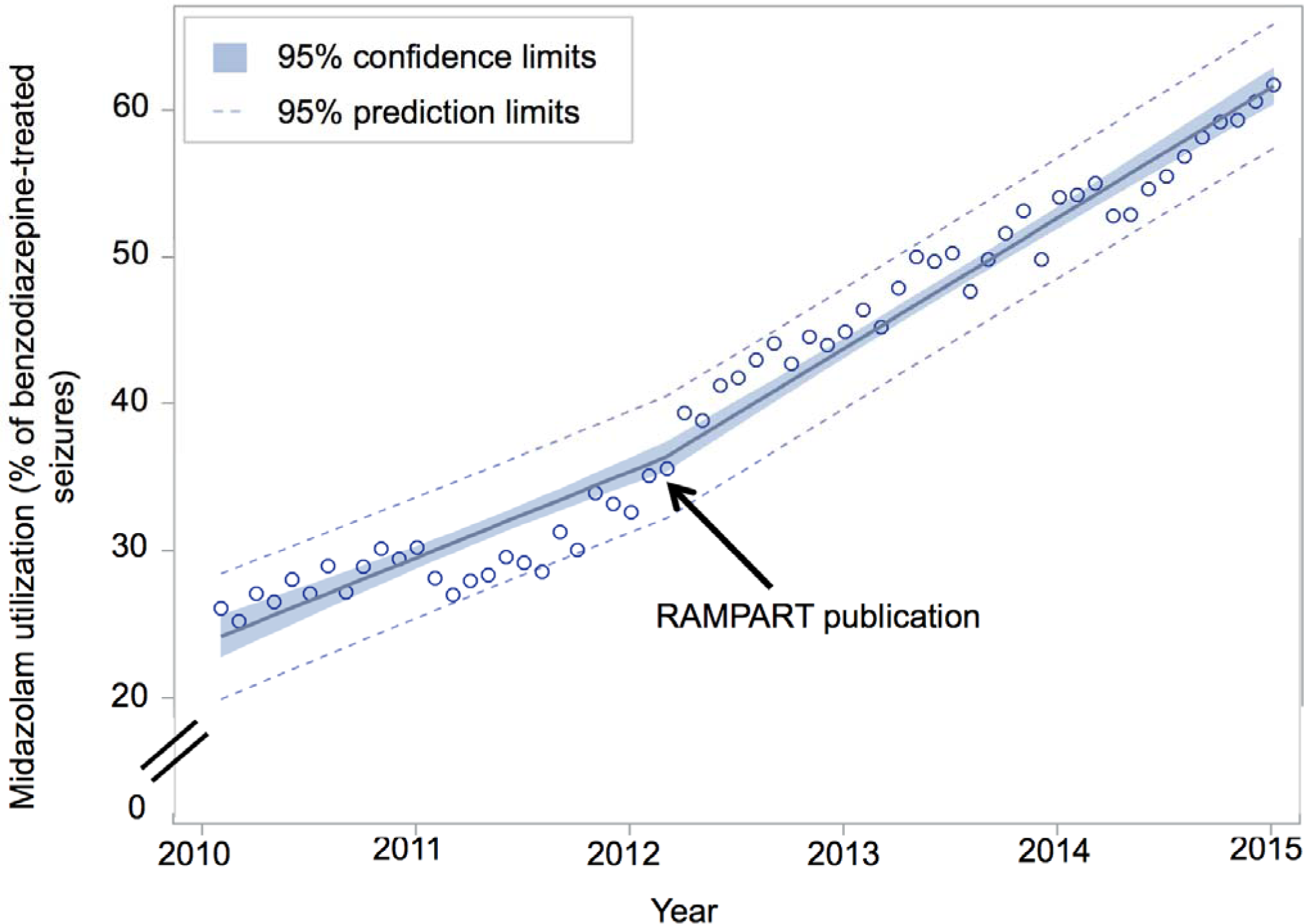
Rate of midazolam adoption before and after RAMPART publication, February 2012. Piecewise linear regression of midazolam utilization among all 3504 EMS agencies in NEMSIS database who treated at least one seizure with benzodiazepines demonstrates a significant increase in the rate of midazolam adoption (pre-RAMPART 5.9% per year; post-RAMPART 8.9% per year; difference +3.0% per year; 95%CI, 1.6%-4.5% per year).

**Fig 4.**
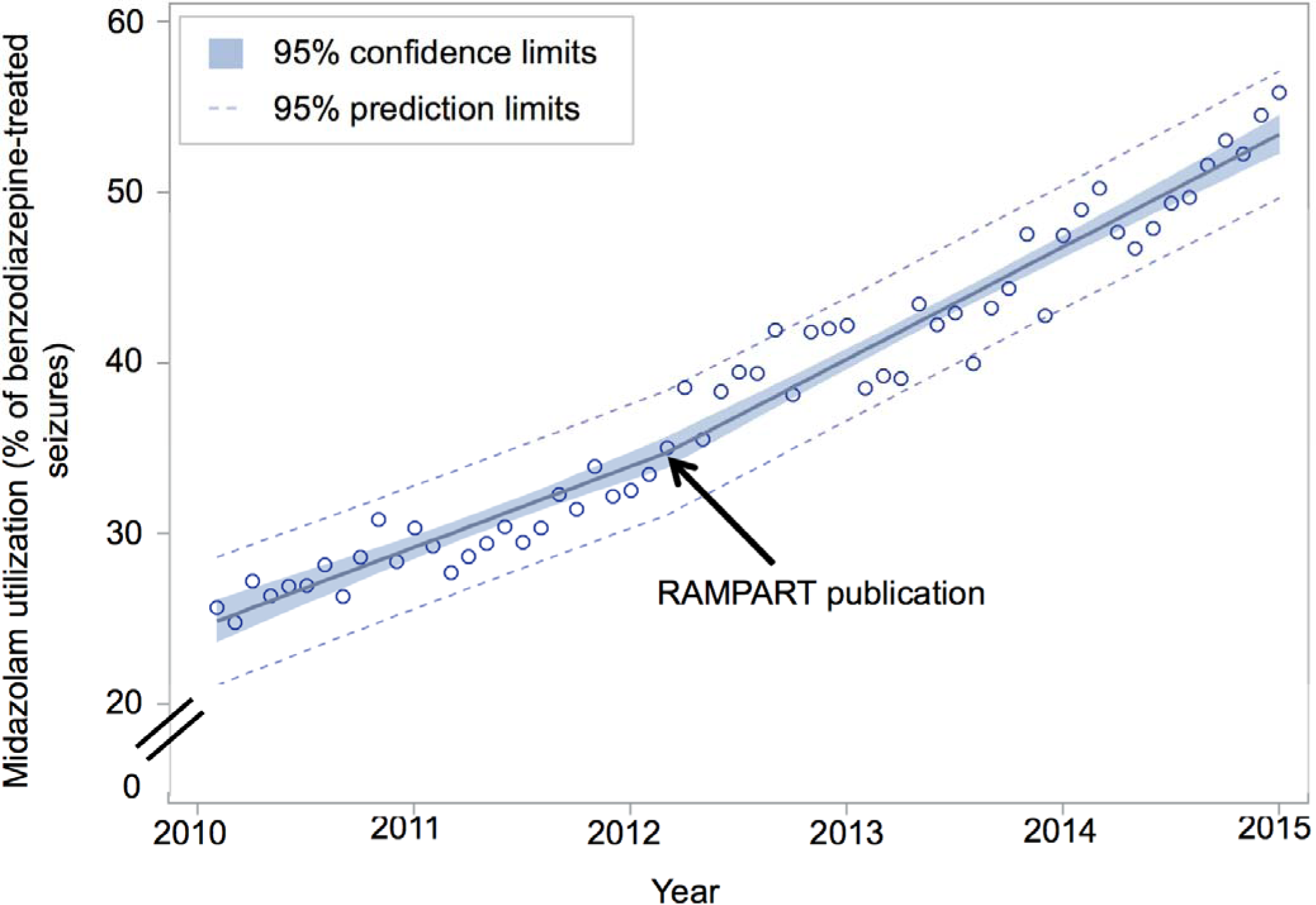
Post-hoc sensitivity analysis of rate of midazolam adoption before and after RAMPART publication among EMS agencies who contributed data to NEMSIS during every study year. Post-hoc piecewise linear regression of midazolam utilization among the 1761 of 3504 (50%) EMS agencies who treated at least one seizure with benzodiazepines, and who contributed data to NEMSIS during all 5 study years demonstrates a significant increase in the rate of midazolam adoption (pre-RAMPART 4.7% per year; post-RAMPART 6.6% per year; difference +1.8% per year; 95%CI, 0.5%-3.1% per year).

**Fig 5.**
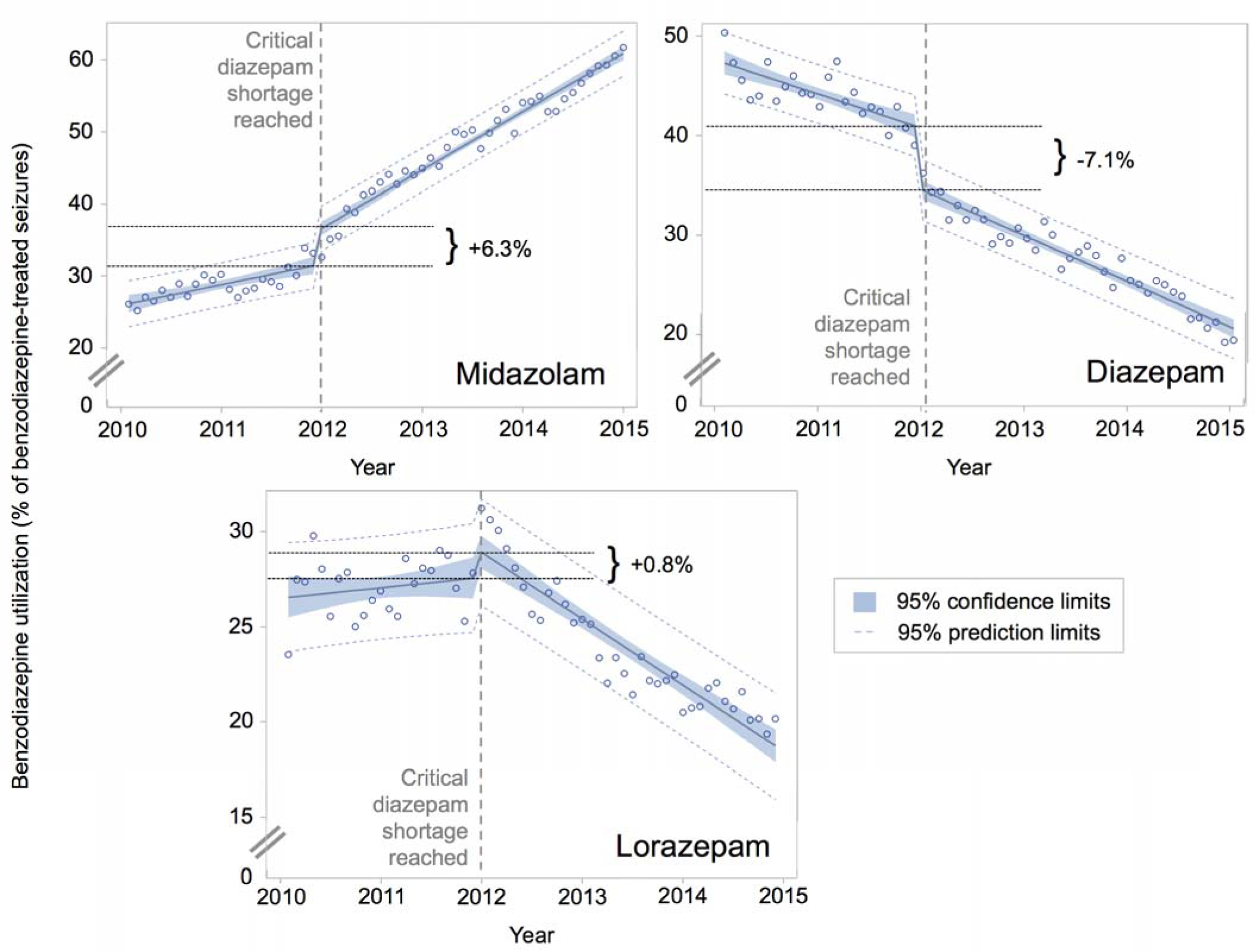
Post-hoc sensitivity analysis of effect of shifts in benzodiazepine use during diazepam shortage on rates of midazolam adoption. Post hoc piecewise linear regression with discontinuous indicator variable for the month in which a critical diazepam shortage was reached. The national diazepam shortage began in August 2011 and in December 2011 became widespread and critical enough to warrant an update to the announcement on ASHP.org that called for use of alternative medications (E. Fox, written communication, May 2016). This call to use alternative benzodiazepines corresponded to the likely depletion of the field inventory of diazepam, which has a typical shelf-life of 2-4 months [6, 7], The resolutions of a prior and the above diazepam shortages, in June 2011 and October 2013, respectively, did not appear to be as impactful on the data (models not shown). This is not surprising as the 2011 shortage affected all Hospira formulations, while in 2013 at least one presentation of diazepam was made available (E. Fox, written communication, May 2016), dampening the impact of the resolution of that shortage. Furthermore, the rolling lorazepam shortage was not modeled, as it was likely buffered by the existence of multiple drug suppliers. The 7.1% decline in diazepam utilization at the start of the shortage was associated with a 6.3% rise in midazolam utilization and a 0.8% rise in lorazepam utilization. After taking into account the sudden shifts in benzodiazepine use that occurred at the time of the shortage, the annual rate of midazolam adoption increased significantly, from 2.9% per year in 2010-2011 to 7.9% per year in 2012-2014 (difference +5.0% per year; 95%CI, 3.0%-7.0% per year).

In the hierarchical logistic regression models, after adjusting for secular trends as well as demographic characteristics (age, gender, region, urbanicity), EMS encounter characteristics (primary role of EMS unit, service type), and agency size (model 4), the odds of receiving midazolam were 24% higher after RAMPART publication (odds ratio [OR] 1.24; 95% CI, 1.17-1.32; Table 3). ICC was high in all models, with 74-76% of variance in midazolam use attributable to the responding EMS agency.

**Table 3:**
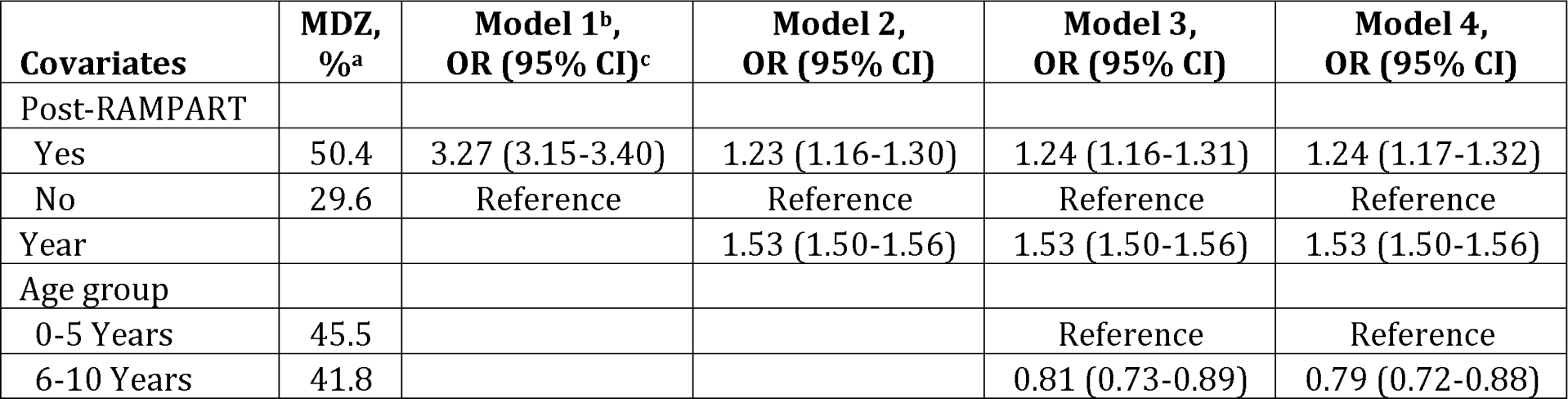
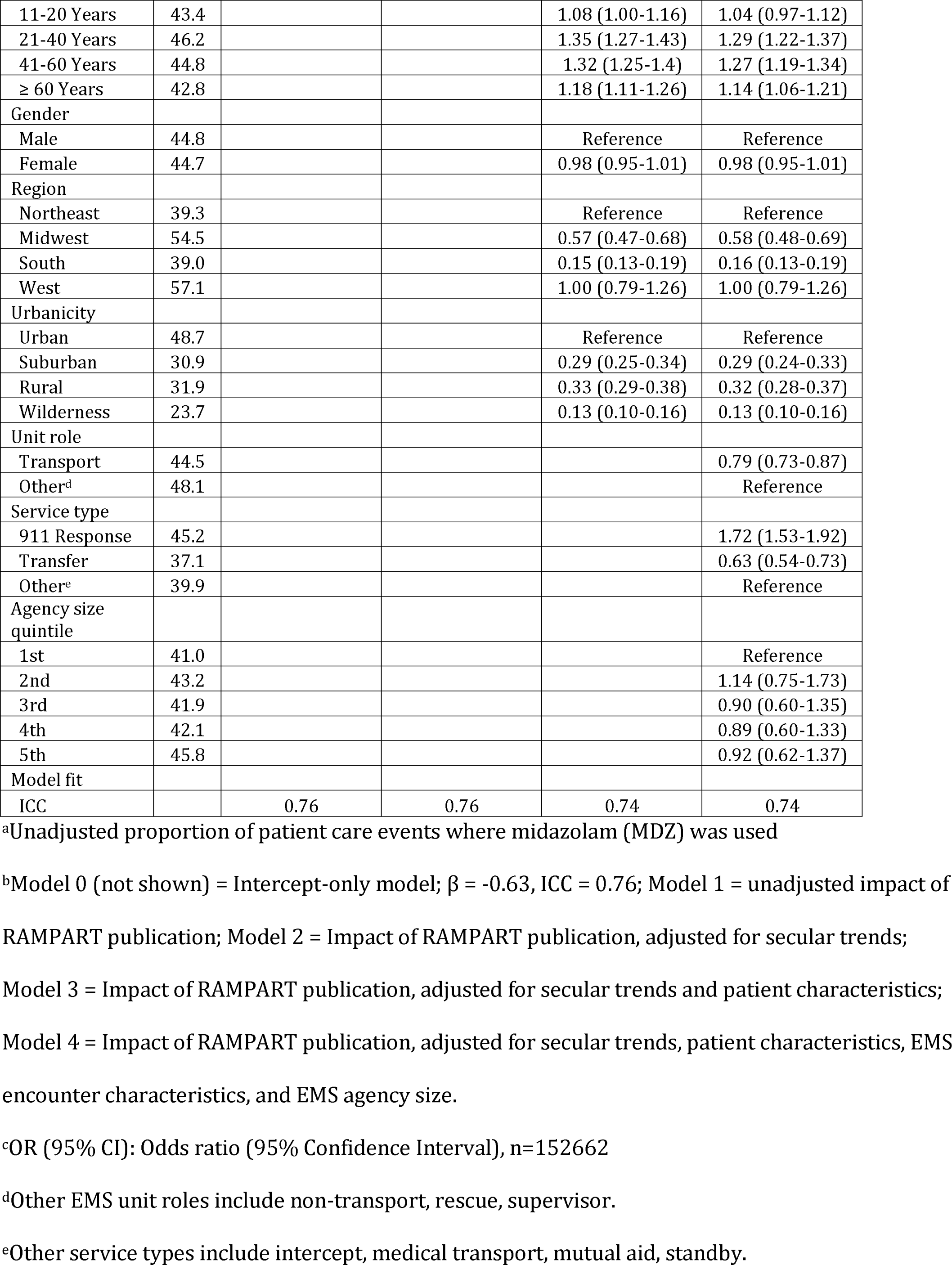
Hierarchical logistic regression models for indicators of midazolam utilization for the treatment of status epilepticus in the pre-hospital setting, 2010-2014.

| Covariates | MDZ, %<sup>a</sup> | Model 1<sup>b</sup>, OR (95% CI)<sup>c</sup> | Model 2, OR (95% CI) | Model 3, OR (95% CI) | Model 4, OR (95% CI) |
| --- | --- | --- | --- | --- | --- |
| Post-RAMPART |  |  |  |  |  |
| Yes | 50.4 | 3.27 (3.15-3.40) | 1.23 (1.16-1.30) | 1.24 (1.16-1.31) | 1.24 (1.17-1.32) |
| No | 29.6 | Reference | Reference | Reference | Reference |
| Year |  |  | 1.53 (1.50-1.56) | 1.53 (1.50-1.56) | 1.53 (1.50-1.56) |
| Age group |  |  |  |  |  |
| 0-5 Years | 45.5 |  |  | Reference | Reference |
| 6-10 Years | 41.8 |  |  | 0.81 (0.73-0.89) | 0.79 (0.72-0.88) |
| 11-20 Years | 43.4 |  |  | 1.08 (1.00-1.16) | 1.04 (0.97-1.12) |
| 21-40 Years | 46.2 |  |  | 1.35 (1.27-1.43) | 1.29 (1.22-1.37) |
| 41-60 Years | 44.8 |  |  | 1.32 (1.25-1.4) | 1.27 (1.19-1.34) |
| ≥ 60 Years | 42.8 |  |  | 1.18 (1.11-1.26) | 1.14 (1.06-1.21) |
| Gender |  |  |  |  |  |
| Male | 44.8 |  |  | Reference | Reference |
| Female | 44.7 |  |  | 0.98 (0.95-1.01) | 0.98 (0.95-1.01) |
| Region |  |  |  |  |  |
| Northeast | 39.3 |  |  | Reference | Reference |
| Midwest | 54.5 |  |  | 0.57 (0.47-0.68) | 0.58 (0.48-0.69) |
| South | 39.0 |  |  | 0.15 (0.13-0.19) | 0.16 (0.13-0.19) |
| West | 57.1 |  |  | 1.00 (0.79-1.26) | 1.00 (0.79-1.26) |
| Urbanicity |  |  |  |  |  |
| Urban | 48.7 |  |  | Reference | Reference |
| Suburban | 30.9 |  |  | 0.29 (0.25-0.34) | 0.29 (0.24-0.33) |
| Rural | 31.9 |  |  | 0.33 (0.29-0.38) | 0.32 (0.28-0.37) |
| Wilderness | 23.7 |  |  | 0.13 (0.10-0.16) | 0.13 (0.10-0.16) |
| Unit role |  |  |  |  |  |
| Transport | 44.5 |  |  |  | 0.79 (0.73-0.87) |
| Other<sup>d</sup> | 48.1 |  |  |  | Reference |
| Service type |  |  |  |  |  |
| 911 Response | 45.2 |  |  |  | 1.72 (1.53-1.92) |
| Transfer | 37.1 |  |  |  | 0.63 (0.54-0.73) |
| Other<sup>e</sup> | 39.9 |  |  |  | Reference |
| Agency size quintile |  |  |  |  |  |
| 1st | 41.0 |  |  |  | Reference |
| 2nd | 43.2 |  |  |  | 1.14 (0.75-1.73) |
| 3rd | 41.9 |  |  |  | 0.90 (0.60-1.35) |
| 4th | 42.1 |  |  |  | 0.89 (0.60-1.33) |
| 5th | 45.8 |  |  |  | 0.92 (0.62-1.37) |
| Model fit |  |  |  |  |  |
| ICC |  | 0.76 | 0.76 | 0.74 | 0.74 |

<sup>a</sup>Unadjusted proportion of patient care events where midazolam (MDZ) was used
<sup>b</sup>Model 0 (not shown) = Intercept-only model; $\beta$ = -0.63, ICC = 0.76; Model 1 = unadjusted impact of RAMPART publication; Model 2 = Impact of RAMPART publication, adjusted for secular trends; Model 3 = Impact of RAMPART publication, adjusted for secular trends and patient characteristics; Model 4 = Impact of RAMPART publication, adjusted for secular trends, patient characteristics, EMS encounter characteristics, and EMS agency size.
<sup>c</sup>OR (95% CI): Odds ratio (95% Confidence Interval), n=152662
<sup>d</sup>Other EMS unit roles include non-transport, rescue, supervisor.
<sup>e</sup>Other service types include intercept, medical transport, mutual aid, standby.

In the fully adjusted model, there was no significant impact of gender on the odds of receiving midazolam. Patients aged 21-60, and those being treated in the Northeast and West, as well as in urban areas, had the highest odds of receiving midazolam. EMS encounter characteristics associated with increased odds of receiving midazolam included treatment by an EMS unit providing 911-scene response or by a unit in a non-transport role. Agency size had no effect on the odds of receiving midazolam. Race, ethnicity, and service level were frequently missing (often by agency and by state) and were not included in the multivariable models.

### Agency-level analysis

Of the 2709 EMS agencies that treated a seizure in more than one year, allowing for comparison between years, 1094 (40%) increased their midazolam use, 476 (18%) decreased use, and 1139 (42%) had no change in use. Of the agencies that did not change their midazolam use, 462 (41%) consistently used midazolam and 632 (55%) never used midazolam.

The results of the post hoc multinomial logistic regression can be seen in Table 4, which represents 7,524 (72%) out of 10,413 agency-years, from agencies that submitted any data to all the explanatory variables.

**Table 4:** Multinomial logistic regression of agency-level variables on odds of midazolam use levels.

| Midazolam use level v. none (n=3153)<sup>a</sup> | Some (n=1595),<br>OR (95% CI)<sup>b</sup> | Most (n=1121),<br>OR (95% CI) | All (n=1655),<br>OR (95% CI) |
| --- | --- | --- | --- |
| Year | 1.21 (1.15-1.27)* | 1.41 (1.34-1.49)* | 1.44 (1.37-1.52)* |
| Patient population<sup>c</sup> |  |  |  |
| 0-5 Year-olds | 1.44 (1.17-1.77)* | 1.40 (1.13-1.74)* | 0.95 (0.75-1.20) |
| 6-20 Year-olds | 1.03 (0.74-1.41) | 1.76 (1.28-2.40)* | 2.61 (2.01-3.38)* |
| ≥ 60 Year-olds | 0.98 (0.89-1.07) | 1.03 (0.93-1.14) | 1.20 (1.10-1.31)* |
| Females | 0.65 (0.56-0.76)* | 0.73 (0.61-0.86)* | 1.01 (0.87-1.16) |
| Blacks | 0.89 (0.85-0.92)* | 0.87 (0.83-0.91)* | 0.89 (0.86-0.93)* |
| Hispanics | 1.16 (1.09-1.23)* | 1.06 (0.99-1.14) | 1.14 (1.07-1.20)* |
| Agency size |  |  |  |
| Average annual call volume<sup>d</sup> | 1.03 (1.03-1.04)* | 1.04 (1.03-1.05)* | 1.02 (1.01-1.02)* |
| Size quintile<sup>e</sup> | 2.65 (2.37-2.96)* | 1.75 (1.57-1.96)* | 0.89 (0.82-0.96)* |
| Catchment area<sup>f</sup> | 1.06 (0.99-1.14) | 1.11 (1.03-1.19)* | 0.94 (0.86-1.02) |
| Urbanicity<sup>g</sup> |  |  |  |
| Urban (n=3769) | Reference | Reference | Reference |
| Suburban (n=631) | 1.09 (0.86-1.39) | 1.27 (0.99-1.64) | 0.51 (0.40-0.65)* |
| Rural (n=1209) | 0.84 (0.69-1.02) | 0.62 (0.49-0.79)* | 0.51 (0.43-0.61)* |
| Wilderness (n=510) | 0.79 (0.58-1.06) | 0.62 (0.44-0.87)* | 0.28 (0.22-0.38)* |
| Mixed (n=1405) | 1.03 (0.84-1.27) | 1.00 (0.80-1.25) | 0.55 (0.43-0.70)* |
| Highest service level |  |  |  |
| BLS (n=267) | Reference | Reference | Reference |
| ALS (n=5586) | 0.97 (0.64-1.47) | 0.50 (0.35-0.73)* | 0.54 (0.39-0.74)* |
| Air (n=1655) | 0.97 (0.63-1.50) | 0.42 (0.28-0.62)* | 0.31 (0.22-0.44)* |
| Specialty care only (n=16) | 4.16 (0.68-25.27) | 1.06 (0.13-8.59) | 2.81 (0.51-15.38) |
| Transport capability |  |  |  |
| Yes (n=7448) | Reference | Reference | Reference |
| No (n=76) | 0.55 (0.26-1.18) | 0.40 (0.15-1.02) | 0.47 (0.25-0.89)* |
| Provides 911 scene response |  |  |  |
| Yes (n=7497) | 0.70 (0.20-2.42) | 1.52 (0.34-6.83) | 1.12 (0.36-3.45) |
| No (n=27) | Reference | Reference | Reference |

<sup>a</sup> n (in agency-years)=7,524, unless otherwise noted for categorical variables
<sup>b</sup> OR (95%CI) = odds ratio (95% confidence interval),\*significant at alpha = 0.05
<sup>c</sup> OR is per 10% increase in proportion of agency's population
<sup>d</sup> OR is per 1000 calls/year increase
<sup>e</sup> OR is per unit increase in size quintile
<sup>f</sup> OR is per unit increase in counties in which the agency operates
<sup>g</sup> Urbanicity level is defined by >95% of an agency's calls in the same level. Mixed urbanicity
includes agencies that had <95% of calls in any one level of urbanicity

### Secondary Outcomes

Overall, independent of agent, frequency of rescue therapy and airway interventions changed little before and after publication of RAMPART. Rescue therapy was used 0.5% (95%CI, 0.05 - 1.0%) more often after publication (10,741 of 42,711, 25.2% pre versus 29,236 of 13,828, 25.7% post), and airway interventions were used 0.3% (95%CI, 0.2 - 0.5%) less often after publication (944 of 42,711, 2.2% pre versus 2134 of 113,828,1.9% post).

Those initially treated with midazolam were both more prevalent in post publication period and had a 3.3% (95%CI, 2.8 - 3.7%) lower rate of use of rescue therapy than those treated with other benzodiazepines (16,632 of 70,045, 23.7% with midazolam versus 23,345 of 86,494, 27.0% with other benzodiazepines). They also had a 1.0% (95%CI, 0.9 - 1.2%) higher rate of airway interventions (1,767 of 70,045, 2.5% versus 1,311 of 86,494,1.5%).

Furthermore, rates of rescue therapy and airway interventions for those treated with midazolam have decreased over time, while the rates associated with treatment with other benzodiazepines have increased or remained constant, respectively (Fig 6). In the fully adjusted hierarchical logistic regression models of secondary outcomes there was a significant interaction between midazolam use and time for the prediction of the odds of airway interventions (p=0.001; Table 5) and rescue therapy (p<0.001; Table 6), with midazolam becoming increasingly safe and effective compared to other benzodiazepines. The paradoxical overall increase in rescue therapy despite midazolam being more effective than other benzodiazepines, and more effective and more prevalent in the post-publication period is likely driven by the increase in rescue therapy over time for other benzodiazepines, as demonstrated by this interaction.

**Fig 6.**
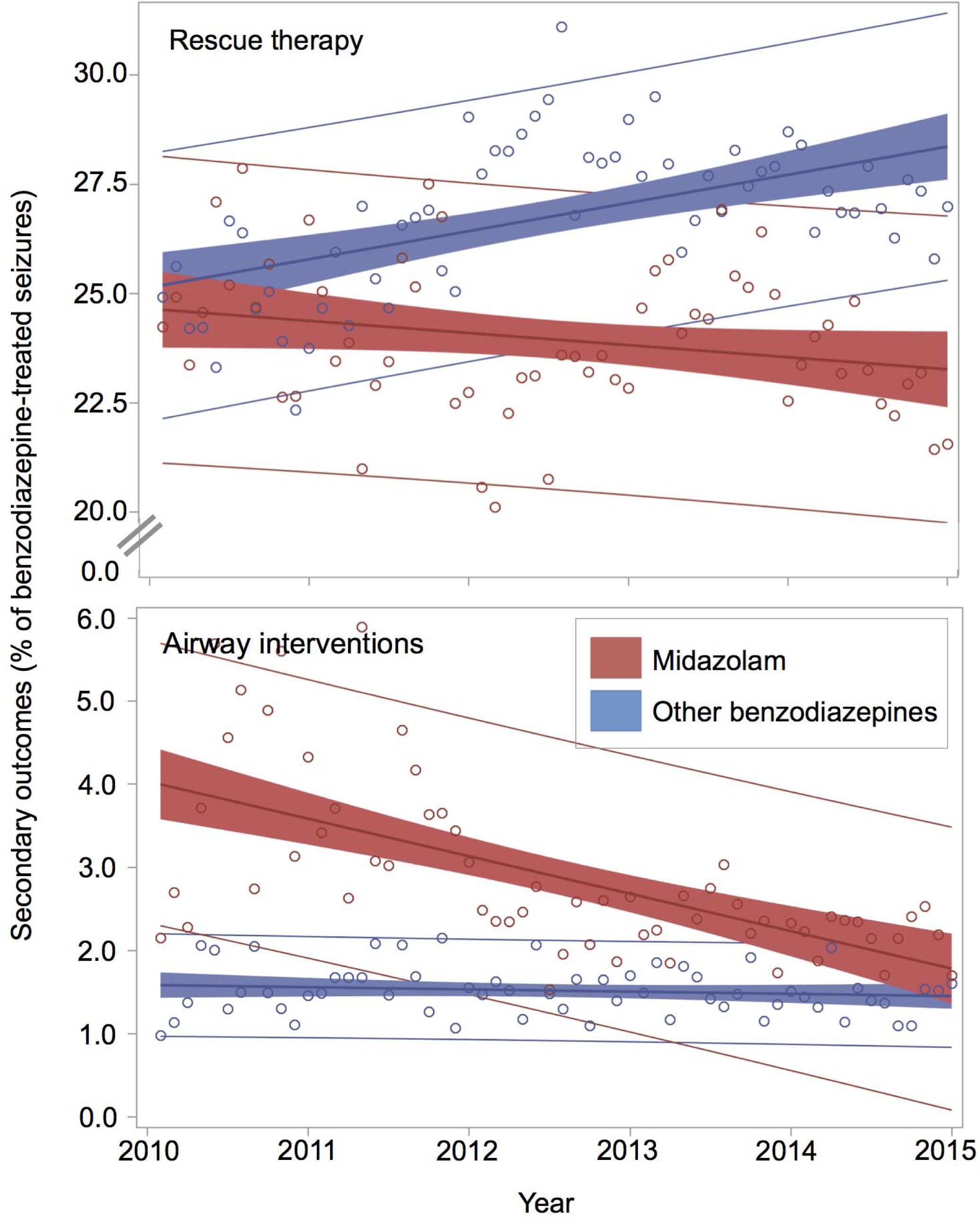
Rates of secondary outcomes for pre-hospital status epilepticus treated with midazolam and other benzodiazepines. There is an interaction between midazolam use and time for the prediction of both secondary outcomes, with midazolam becoming increasingly safe and effective relative to other benzodiazepines.

**Table 5:**
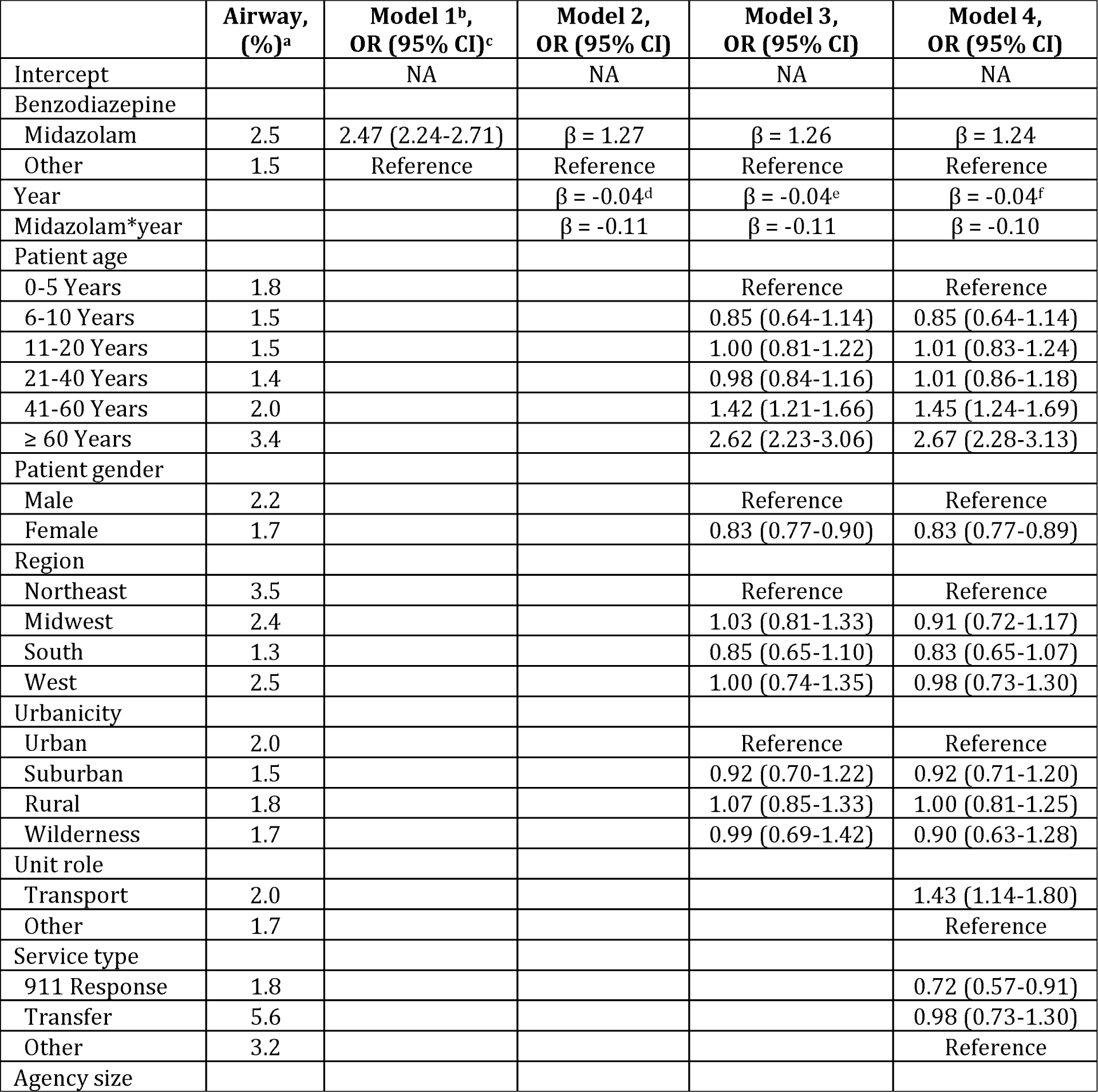
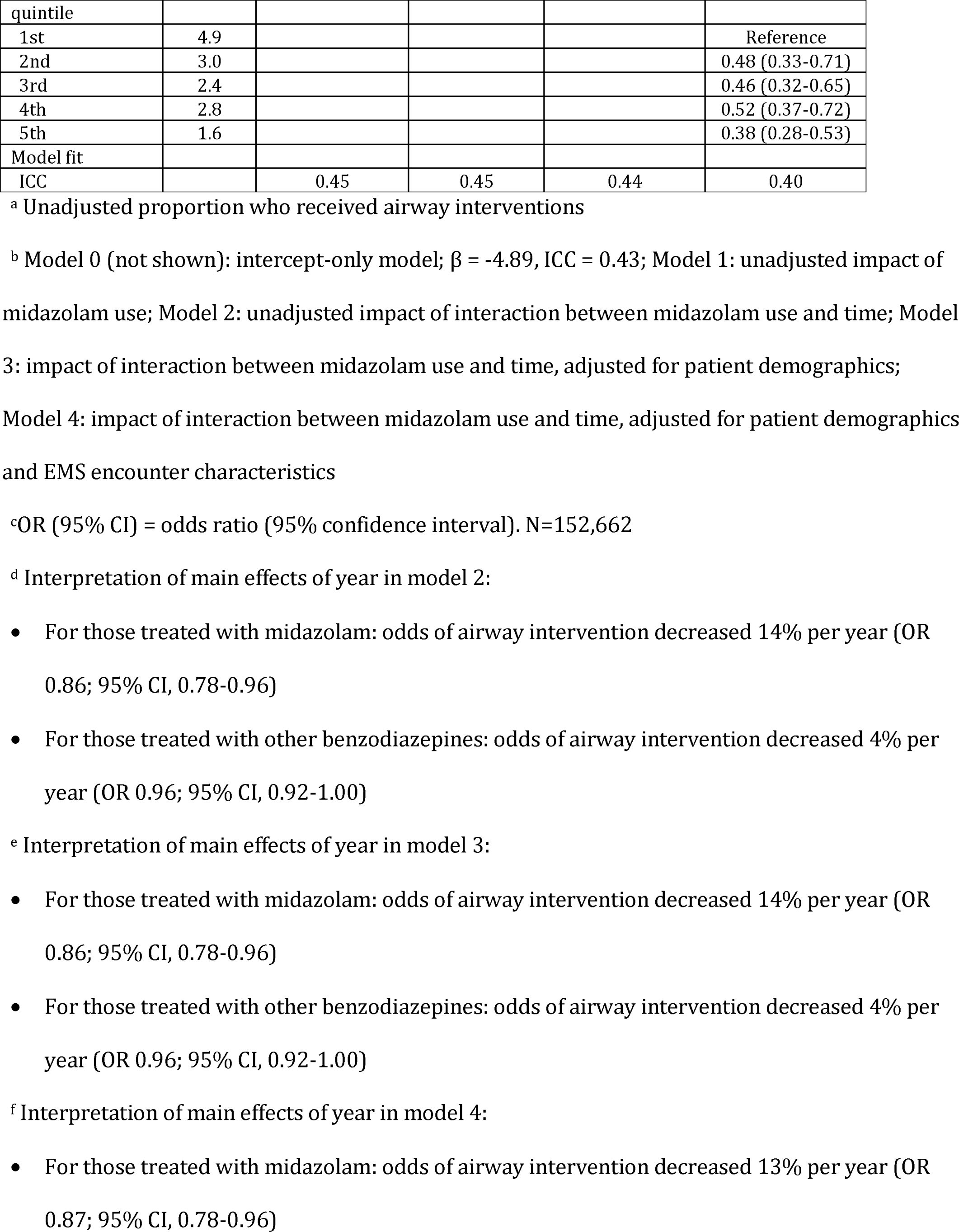
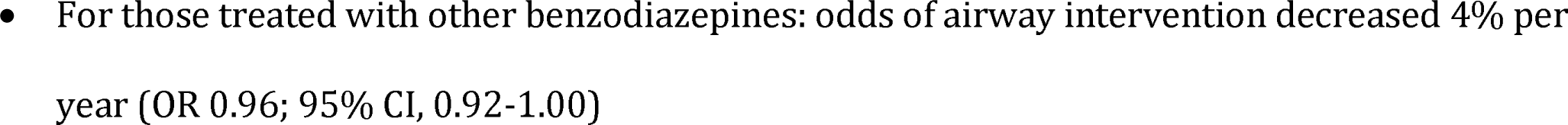
Hierarchical logistic regression models for indicators of airway interventions among patients with status epilepticus in the pre-hospital setting.

**Table 6:** Hierarchical logistic regression models for indicators of rescue therapy after initial treatment of status epilepticus with benzodiazepines in the pre-hospital setting.

|  | Rescue,<br>(%)<sup>a</sup> | Model 1<sup>b</sup>,<br>OR (95% CI)<sup>c</sup> | Model 2,<br>OR (95% CI) | Model 3,<br>OR (95% CI) | Model 4,<br>OR (95% CI) |
| --- | --- | --- | --- | --- | --- |
| Intercept |  | NA | NA | NA | NA |
| Benzodiazepine |  |  |  |  |  |
| Midazolam | 23.7 | 0.83 (0.81-0.86) | $\beta = -0.02$ | $\beta = -0.06$ | $\beta = -0.06$ |
| Other | 27.0 | Reference | Reference | Reference | Reference |
| Year | | | $\beta = 0.03^d$ | $\beta = 0.03^e$ | $\beta = 0.03^f$ |
| Midazolam*year | | | $\beta = -0.06$ | $\beta = -0.05$ | $\beta = -0.05$ |
| Patient age |  |  |  |  |  |
| 0-5 Years | 22.2 |  |  | Reference | Reference |
| 6-10 Years | 21.9 |  |  | 1.01 (0.93-1.10) | 1.03 (0.94-1.12) |
| 11-20 Years | 26.5 |  |  | 1.32 (1.25-1.41) | 1.37 (1.29-1.45) |
| 21-40 Years | 28.1 |  |  | 1.48 (1.41-1.56) | 1.55 (1.47-1.62) |
| 41-60 Years | 25.1 |  |  | 1.27 (1.21-1.34) | 1.32 (1.26-1.39) |
| ≥ 60 Years | 23.4 |  |  | 1.14 (1.08-1.20) | 1.19 (1.12-1.25) |
| Patient gender |  |  |  |  |  |
| Male | 25.7 |  |  | Reference | Reference |
| Female | 25.4 |  |  | 0.96 (0.94-0.99) | 0.96 (0.94-0.99) |
| Region |  |  |  |  |  |
| Northeast | 29.3 |  |  | Reference | Reference |
| Midwest | 28.1 |  |  | 1.01 (0.92-1.11) | 0.94 (0.86-1.03) |
| South | 21.7 |  |  | 0.53 (0.49-0.58) | 0.53 (0.49-0.58) |
| West | 32.5 |  |  | 1.03 (0.92-1.15) | 1.01 (0.91-1.12) |
| Urbanicity |  |  |  |  |  |
| Urban | 25.6 |  |  | Reference | Reference |
| Suburban | 22.0 |  |  | 0.81 (0.74-0.89) | 0.80 (0.74-0.88) |
| Rural | 27.1 |  |  | 0.96 (0.89-1.04) | 0.94 (0.87-1.01) |
| Wilderness | 27.3 |  |  | 0.90 (0.80-1.02) | 0.85 (0.75-0.96) |
| Unit role |  |  |  |  |  |
| Transport | 25.7 |  |  |  | 1.18 (1.09-1.26) |
| Other | 24.2 |  |  |  | Reference |
| Service type |  |  |  |  |  |
| 911 Response | 24.7 |  |  |  | 0.89 (0.81-0.97) |
| Transfer | 43.5 |  |  |  | 2.07 (1.85-2.32) |
| Other | 27.7 |  |  |  | Reference |
| Agency size<br>quintile |  |  |  |  |  |
| 1st | 37.2 |  |  |  | Reference |
| 2nd | 32.9 |  |  |  | 0.84 (0.73-0.97) |
| 3rd | 31.7 |  |  |  | 0.80 (0.70-0.92) |
| 4th | 29.6 |  |  |  | 0.76 (0.67-0.87) |
| 5th | 23.5 |  |  |  | 0.64 (0.56-0.73) |
| Model fit |  |  |  |  |  |
| ICC |  | 0.17 | 0.17 | 0.15 | 0.13 |

<sup>a</sup> Unadjusted proportion who received rescue therapy
<sup>b</sup> Model 0 (not shown): intercept-only model; $\beta = -1.11$ , ICC = 0.17; Model 1: unadjusted impact of midazolam use; Model 2: unadjusted impact of interaction between midazolam use and time; Model 3: impact of interaction between midazolam use and time, adjusted for patient demographics; Model 4: impact of interaction between midazolam use and time, adjusted for patient demographics and EMS encounter characteristics
<sup>c</sup> OR (95% CI) = odds ratio (95% confidence interval). N=152,662
<sup>c</sup> Interpretation of main effects of year in model 2:
- 403 • For those treated with midazolam: odds of rescue therapy decreased 2% per year (OR 0.98; 95% CI, 0.95-1.01) - 405 • For those treated with other benzodiazepines: odds of rescue therapy increased 3% per year (OR 1.03; 95% CI, 1.02-1.05)
<sup>e</sup> Interpretation of main effects of year in model 3:
- 408 • For those treated with midazolam: odds of rescue therapy decreased 2% per year (OR 0.98; 95% CI, 0.95-1.01) - 410 • For those treated with other benzodiazepines: odds of rescue therapy increased 3% per year (OR 1.03; 95% CI, 1.02-1.05)
<sup>f</sup> Interpretation of main effects of year in model 4:
- 413 • For those treated with midazolam: odds of rescue therapy decreased 2% per year (OR 0.98; 95% CI, 0.95-1.02) - 415 • For those treated with other benzodiazepines: odds of rescue therapy increased 3% per year (OR 1.03; 95% CI, 1.02-1.04)

## Discussion

In this retrospective, observational, cohort study, we found that midazolam use for the prehospital treatment of status epilepticus increased substantially from 2010-2014, and the rate of adoption increased significantly after the publication of RAMPART. These data are consistent with effective but incomplete clinical translation of the RAMPART results, but the effects of the trial cannot be isolated. The nature of underlying trends is unclear and the potential effects of already changing practices, drug shortages, and early release of trial results to the Federal Drug Administration and the EMS agencies that participated in RAMPART are potentially confounding. Increasing use of midazolam in the pre-publication phase may represent a pre-existing trend that is entirely agnostic to the trial. If so, this trend may or may not have continued in the post-publication period even if there had been no trial data. In this scenario, the trial data may have had little effect on the secular trend, may have been permissive, or may have positively reinforced the trend. It is also possible that the trend in the pre-publication period may be related to the occurrence of the trial, reducing the apparent impact of the published results. The relatively steep pre-publication trend in midazolam use is consistent with this later hypothesis, as a 5.9% per year rate of increase is not likely to have been sustained for more than 4 years prior to the study period. However, the nature of the trend prior to the study period is unknown, as is its association with other publications on midazolam’s efficacy[7]. These observational data also confirm real world efficacy and safety findings identified in the well-controlled clinical trial, thereby supporting the use of midazolam by EMS personnel.

After adjusting for secular trends and other factors, the odds of receiving midazolam were 24% higher after the publication of RAMPART, which reflects the significantly increased rate of adoption of midazolam around that time. The increasing use of midazolam could be explained by growing acknowledgement of the benefits of midazolam for the treatment of pre-hospital status epilepticus. Midazolam is not only as effective as alternative intravenous medications [3, 5], but is also easier to administer[8], has better pharmacodynamics than diazepam [9], is cheaper [10-12] and has a longer shelf life than lorazepam [13]. Furthermore, midazolam is subject to far fewer shortages, with diazepam and lorazepam making up 2 of the 10 most susceptible drugs to national shortages [14]. While our results suggest that the diazepam shortage did not by-itself drive the increased rate of midazolam adoption, it may have contributed. Certainly, this trend could partially result from growing frustration with repeated shortages of alternative benzodiazepines.

The increasing use of midazolam relative to alternative benzodiazepines is consistent with the effects of several alternative or additive events, as described above, including the effective but incomplete clinical translation of the RAMPART results. The problem of knowledge translation has been distilled into two distinct components: “getting the evidence straight” and “getting the evidence used” [2]. There has been a growing body of literature supporting the use of midazolam for the treatment of status epilepticus [3, 7,8,15], and this study suggests that the increase in midazolam use could be related to this growing consensus on midazolam’s safety and efficacy. The as-of-yet incomplete penetration of midazolam is not surprising in light of the tremendous amount of time it typically takes for new knowledge generated by randomized controlled trials to be broadly incorporated into practice [1]. Even where evidence has been integrated into clinical guidelines, dissemination and implementation of those guidelines in pre-hospital clinical practice is often slow and cannot be said to be automatic [16].

There were small but statistically significant differences in rates of rescue therapy and airway interventions before and after RAMPART publication and between seizures treated with midazolam and other benzodiazepines. Given that the magnitude of these differences are small and their directions opposite, the increased use of midazolam was not associated with increased complications overall. Furthermore, the safety and efficacy of midazolam improved over time relative to other benzodiazepines. These results likely reflect changing patterns of midazolam dose, route of administration, or both. Since the NEMSIS database does not have a reliable indicator for the dose or route of administration, we were unable to test this speculation or the possible impact of RAMPART on these patterns of use. Interestingly, if the increased efficacy over time reflects an increase in midazolam dose, then the decreased intubation rate with midazolam over time might suggest that airway intervention is more likely to be a ramification of treatment failure rather than of over-sedation—a relationship that has been previously observed but that is still not well defined [17]. Alternatively, if our data reflect changes in the route of midazolam administration, our study could indicate differences in safety and efficacy between, for example, the intramuscular and intranasal route.

## Limitations

Several limitations of this study should be noted. The study utilized data that were collected by EMS personnel for purposes other than this study and therefore have limited ability to answer all study questions. Additional limitations of the NEMSIS dataset have been described elsewhere [4]. Notably, NEMSIS is not a population-based dataset, which limits the ability to draw conclusions about relative risks of treatment with midazolam versus other benzodiazepines in the general population.

We found that the overall intubation rate for all benzodiazepine-treated seizures was 2.0%, which is far less than other reports [15,17–19]. This discrepancy likely indicates that our operational definition of status epilepticus as any benzodiazepine-treated seizure is too broad, representing a far more encompassing, less severely ill, patient population. The overtreatment of seizures that do not meet criteria for status epilepticus could introduce bias if there are significant differences in the benzodiazepine used for seizures of different acuity levels—a variable we were unable to assess.

## Conclusions

Our data are consistent with effective, ongoing, but incomplete clinical translation of the RAMPART results. The effects of the trial, however, could not be isolated. The safety and effectiveness of midazolam for treatment of seizures in the prehospital setting in clinical practice appear consistent with trial data. Along with midazolam’s ease of administration, low cost, long shelf life, and low susceptibility to shortages, this should encourage continuing increases in midazolam utilization. Since the use of midazolam is highly dependent on the responding EMS agency, future research should focus on the decision-making process of EMS medical directors across the country.

## Acknowledgements

We would like to acknowledge Henry Wang M.D. M.S., Clay Mann Ph.D. M.S., Mengtao Dai M.S., and Karen Jacobson B.A. NREMT-P from the NEMSIS Technical Assistance Center for their assistance in the preparation of the database; Eric Lavonas M.D. for his description of a concurrent diazepam shortage within our study period; Steve Leber M.D., Ph.D., and Stephanie Chalifour B.A., for their editing assistance in review of the manuscript. No compensation was received for the above contributions.

